# 5-iodotubercidin sensitizes cells to RIPK1-dependent necroptosis by interfering with NFκB signaling

**DOI:** 10.1101/2023.03.03.530727

**Authors:** Chanchal Chauhan, Andreas Kraemer, Stefan Knapp, Mark Windheim, Alexey Kotlyarov, Manoj B. Menon, Matthias Gaestel

**Affiliations:** Institute of Cell Biochemistry, Hannover Medical School, Hannover, 30625, Germany; Institute of Pharmaceutical Chemistry, Goethe University Frankfurt am Main, 60438, Frankfurt am Main, Germany; Structural Genomics Consortium (SGC), Buchmann Institute for Life sciences (BMLS), Goethe University Frankfurt am Main, 60438, Frankfurt am Main, Germany; Frankfurt Cancer Institute (FCI) and German translational cancer network (DKTK) site Frankfurt-Mainz, 60438, Frankfurt am Main, Germany; Kusuma School of Biological Sciences, Indian Institute of Technology Delhi, New Delhi, 110016, India

## Abstract

Receptor-interacting protein kinases (RIPK) −1 and −3 are master regulators of cell fate decisions in response to diverse stimuli and are subjected to multiple checkpoint controls. Earlier studies have established the presence of distinct IKK1/2 and p38/MK2-dependent checkpoints which suppress RIPK1 activation by directly phosphorylating it at different residues. In the present study, we investigated TNF-induced death in MAPK-activated protein kinase 2 (MK2)-deficient cells and show that MK2-deficiency or inactivation predominantly results in necroptotic cell death, even in the absence of caspase inhibition. While MK2-deficient cells can be rescued from necroptosis by RIPK1 inhibitors, RIPK3 inhibition seems to revert the process triggering apoptosis. To understand the mechanism of this necroptosis switch, we screened a 149-compound kinase inhibitor library for compounds which preferentially sensitize MK2-deficient MEFs to TNF-induced cell death. The most potent inhibitor identified was 5-Iodotubericidin, an adenosine analogue acting as adenosine kinase and protein kinase inhibitor. 5-ITu also potentiated LPS-induced necroptosis when combined with MK2 inhibition in RAW264.7 macrophages. Further mechanistic studies revealed that 5-Iodotubericidin induces RIPK1-dependent necroptosis in the absence of MK2 activity by suppressing IKK signaling. The identification of this role for the multitarget kinase inhibitor 5-ITu in TNF-, LPS- and chemotherapeutics-induced necroptosis will have potential implications in RIPK1-targeted therapies.

## Introduction

The TNF-receptor-interacting protein kinases (RIPK) −1 and −3 regulate cell fate in response to diverse stimuli and are subjected to multiple checkpoint controls (reviewed in Humphries et al., 2015; Varfolomeev et al., 2022). RIPK1 is recruited to the TNF receptor and related death-receptors and modulates pro-survival gene expression, inflammation and cell death signaling. Receptor ligandbinding induces subsequent ubiquitination of RIPK1, promoting formation of the complex I, which predominantly modulates pro-survival signaling (Micheau et al., 2003). Receptor-associated RIPK1 in complex I is required for the activation of MAP kinases and NFκB-mediated pro-survival gene expression. Activated RIPK1 may subsequently be released from the receptor to form a cytosolic complex containing CASP8 and FADD to mediate apoptotic cell death (complex IIb) (Wang et al., 2008, Micheau et al., 2003). In the absence of CASP8, RIPK3 can be recruited to RIPK1 inducing subsequent MLKL phosphorylation, oligomerization and necroptosis (Cho et al., 2009; Sun et al., 2012). RIPK1 is also involved in cell death and inflammatory signaling downstream to the interferon-alpha (IFNαR) (McComb et al., 2014) and Toll-like receptors (TLR)-3 and −4 (Meylan et al., 2004, He et al., 2011). Interestingly, treatment of myeloid cells with TLR4-activating bacterial lipopolysaccharide (LPS) in combination with the pan-caspase inhibitor zVAD-fmk (zVAD) results in RIPK1-dependent necroptosis (He et al., 2011). In response to genotoxic stress and depletion of inhibitors of apoptosis proteins (IAPs), RIPK1 also assembles into the ripoptosome, a high molecular weight cytotoxic protein complex, inducing cell death independent of receptor ligation (Tenev et al., 2011).

Therefore, RIPK1 is at the center of a signaling hub, which determines the outcome of cell fate decisions in the context of TNF, TLR and genotoxic signaling. A complex array of posttranslational modifications (PTMs) of RIPK1 determines the balance between cell death and pro-survival signaling (reviewed in (Varfolomeev et al., 2022; Annibaldi et al., 2018)). Receptor-associated ubiquitination of RIPK1 is essential for the recruiting MAP3K7/TAK1 and the IKKα/ß/y resulting in MAPK and NFκB pathway activation, respectively. Deubiquitinated RIPK1 undergoes autophosphorylation at S166 in its activation loop being the best characterized autophosphorylation site associated with kinase activation. Diverse protein kinases have been implicated in phosphorylating RIPK1 and regulating the switch from pro-survival signaling to kinase activation and subsequent assembly of cytotoxic complexes in the cytoplasm including ripoptosome and/or complex IIb and necrosome. Independent of its canonical role in activation of pro-survival NFκB signaling, the inhibitor of κB (IκB)-kinases IKKα/IKK1 and IKKβ/IKK2 have been demonstrated to phosphorylate RIPK1 in complex I, suppressing RIPK1 activation and assembly of complex IIb (Dondelinger et al., 2015). Furthermore, MAPK-activated protein kinase 2 (MK2)-mediated cytoplasmic phosphorylation of RIPK1 at serines S321 and S336 acts as a second checkpoint which suppresses both, RIPK1-S166 autophosphorylation and the assembly of complex IIb (Menon et al., 2017; Dondelinger et al., 2017; Jaco et al., 2017). In addition to IKK1/2 (S25) (Dondelinger et al., 2019) and MK2 (S320, S335) (Menon et al., 2017), it has been recently shown that also TBK1/IKKε (T189) (Lafont et al., 2018) and JAK1/SRC (Y384) (Tu et al., 2022) phosphorylate RIPK1, resulting in suppression of the cytotoxic response. Moreover, recent evidence indicates a role for PPP1R3G/PP1γ mediated dephosphorylation in counteracting kinase-mediated suppression of RIPK1-dependent apoptosis and necroptosis (Du et al., 2021). While the mechanisms of modulation of the intra- and intermolecular interplay and S166 autophosphorylation by these PTMs are still unknown, it became obvious that interfering with these RIPK1-kinases by small molecule inhibitors sensitizes cells to TNF-induced cell death. While single loss of MK2 does not have a strong impact on enhancing TNF-induced cytotoxicity, its combination with IKK inhibitors revealed an MK2-dependent checkpoint in LPS-zVAD-induced necroptosis (Menon et al., 2017). In addition, p38/MK2 inhibition facilitates smac-mimetics (SM)-induced ripoptosome assembly and autocrine TNF production in myeloid cells and has been proposed as a means of circumventing SM-resistance in leukemia (Lalaoui et al., 2016; Jaco et al., 2017; Rijal et al., 2018).

5-Iodotubercidin (5-ITu) is a halogenated pyrrolopyrimidine analogue and this nucleoside analogue has been reported as a potent inhibitor of adenosine uptake and adenosine kinase activity (Davies et al., 1984). While originally used as an adenosine kinase inhibitor, later studies revealed its potential as a pan-protein kinase inhibitor (Massillon et al., 1994). 5-ITu was shown to enhance human β-cell proliferation and glucose-dependent insulin secretion by targeting DYRK1A kinase (Dirice et al., 2016). Low concentration of 5-ITu also inhibits Haspin/GSG2, a histone-H3 kinase (Antoni et al., 2012; Karanika et al., 2020; Heroven et al., 2018). It has also been described as a potent chemotherapeutic agent with strong antitumor activity (Zhang et al., 2013). Recently, 5-ITu has been shown to inhibit SARS-CoV2 replication, via an adenosine kinase-dependent mechanism (Zhao et al., 2022).

The switch between pro-survival and cytotoxic functions of RIPK1 is regulated by diverse post-translational modifications acting as checkpoints for RIPK1 activation (Wang et al., 2021). To understand the MK2-dependent checkpoint on RIPK1 autophosphorylation and the assembly mechanisms of complex IIb and necrosome, we analysed the mode of cell death in MK2-deficient cells. Interestingly, MK2-deficiency is not only associated with enhanced RIPK1 activation, but also exerts a predominantly necroptotic mode of cell death. In a screen for small molecule inhibitors to specifically sensitize MK2-deficient cells to TNF-induced necroptosis, we identified 5-ITu as a prominent regulator of RIPK1-dependent cell death. We present evidence that 5-ITu acts as a modulator of the NFκB pathway and, therefore, compromises the IKK-dependent checkpoint of RIPK1 activation. Our findings indicate that 5-ITu- and MK2-inhibition could have additive effects in sensitizing cells to RIPK1-dependent necroptosis.

## Materials and Methods

### Cell culture

SV40-T antigen immortalized WT, MK2- and MK2/3-deficient (DKO) mouse embryonic fibroblasts (MEFs) and retroviral rescued cells were reported previously (Ronkina et al., 2007; Ronkina et al., 2011). MEFs and RAW 264.7 cells were cultured in DMEM supplemented with 10% fetal calf serum (FCS) and 1% Penicillin/Streptomycin at 37□°C, in a humidified atmosphere supplemented with 5% CO_2_.□

### Antibodies and reagents

MLKL (#14993), Caspase-8 (#4927), p38 MAPK (#9212), MAPKAPK2 (D1E11) (#12155), pS166-RIPK1 (#65746), pS345-MLKL (#37333), phospho-p44/42 MAPK (Erk1/2) (Thr202/Tyr204) (#4370), phospho-p38 MAPK (Thr180/Tyr182) (#9215), phospho-SAPK/JNK MAPK (Thr183/Tyr185) (G9) (#9255), SAPK/JNK (#9252), pS32/36-IkBα (#9246), IkBα (#9242), pS176/180-IKKα/β (#2697), IKKα (#2682), Cleaved-Asp387-Caspase-8 (#8592) antibodies were from Cell Signaling Technology (CST). Further antibodies used were against RIPK1 (#610459, BD Biosciences), GAPDH (#MAB374, Millipore), EF2 (sc-166415, Santa Cruz Biotechnology), ERK2 (sc-154, Santa Cruz Biotechnology). The secondary antibodies used were goat anti-mouse IgG (H+L)-HRP (#115-035-003, Dianova) and goat anti-rabbit IgG (H+L)-HRP (111-035-003, Dianova).

### Kinase inhibitors

The kinase inhibitor library, composed of 149 selective or broad-spectrum kinase inhibitors dissolved in DMSO (10 mM stock concentration), was purchased from Cayman chemicals (#10505). The following reagents were used for cell treatments at given concentrations: Lipopolysaccharide (*Escherichia coli* O55:B5, #L2880, Sigma-Aldrich, 100□ng□ml^-1^), recombinant human TNFα (rHuTNF, #50435.50, Biomol, 10□ng/ml), Birinapant/Smac Mimetics (#HY-16591, MedChem Express, 1μM), pan-caspase inhibitor zVAD-fmk (#4026865.0005, Bachem, 25□μM), MK2 inhibitor PF3644022 (#4279, Tocris, 5μM), RIPK1 inhibitor Nec-1 (#BML-AP309-0020, Enzo Life Sciences, 50μM), RIPK1 inhibitor Nec-1s (# 10-4544-5mg, Tebu-Bio, 50μM), RIPK3 inhibitor GSK872 (#HY-101872, MedChem Express, 5μM), IKKβ inhibitor BMS345541 (#Axon 1731, Axon Medchem, 5μM), TBK1/IKKε inhibitor BX795 (#T1830, Tebu-Bio), 5-Iodotubericidin (#HY-15424, MedChem Express, 2.5-10μM), ABT-702 dihydrochloride (#HY-103161, MedChem Express, 5μM), Staurosporine (#81590, Cayman, 5μM), Etoposide (#E1383, Sigma, 5μM), Doxorubicin (#15007, Cayman, 5μM), Gemcitabine (#S1714, Selleckchem, 100nM)

### Western blotting

Cells were lysed directly in cell culture plates with 2x SDS sample buffer containing 10% SDS, 1.5M TRIS pH 8.8, 5% Glycerol, 2.5% 2-Mercaptoethanol, and Bromophenol Blue, followed by denaturing the samples for 5 minutes at 95°C. SDS-PAGE (7.5–16% gradient) gels were used for protein separation followed by western blotting using nitrocellulose membrane (#10600001, Amersham Biosciences). Blotted membranes were blocked with 5% powdered skim milk in PBS with 0.1% Tween 20 for 1h at room temperature. PBS with 0.1% Tween 20 was used for washing the membranes three times followed by incubation with the primary antibody overnight, at 4 °C. Next day, blots were washed and incubated with horseradish peroxidase-conjugated secondary antibodies for one hour, at room temperature. ECL detection kit, WESTAR NOVA 2.0 (#XLS07105007, 7Biosciences) was used for the detection of proteins and the digital chemiluminescence images were taken by a Luminescent Image Analyser LAS-3000 (Fujifilm).

### Analysis of cell viability and cell death

Cell Counting Kit-8 (CCK-8) assay (Bimake, #B334304) was used to measure cell viability. MEF cells and RAW264.7 cells were seeded in triplicates in a 96-well format in 100μl media per well. Next day, cells were treated with various indicated combinations of reagents. For MEFs, the medium was changed with diluted WST-8 for viability estimation. For RAW 264.7 cells, 10μl WST-8 was added directly to the wells in complete medium. The plates were then incubated in a cell culture incubator for 1 hour at 37°C until the colour turned orange and the absorbance was measured at 450nm using a microplate reader (Perkin Elmer Wallac Victor2 1420 Multilabel Counter). Results are expressed as the percentage of cell viability per well in relation to the maximum cell viability of DMSO-treated cells. For inhibitor screen, MEFs were seeded in triplicates and treated with 10 μM inhibitor samples of the panel, alone or in combination with 10 ng/ml TNF. For measuring cell death, the cell impermeable Sytox Green Nucleic Acid Stain (Invitrogen) was added to the samples after 30 minutes of TNF treatment, at a final concentration of 0.25μM. The indicated treatments were performed in triplicates in a 96-well plate format. A microplate reader (Perkin Elmer Wallac Victor2 1420 Multilabel Counter) was used to measure the kinetics of fluorescence.

### Differential scanning fluorimetry (DSF)-based selectivity screening against a curated kinase IIbrary

The assay was performed as previously described (Krämer et al., 2020; Federov et al., 2012). Briefly, recombinant protein kinase domains at a concentration of 20μM were mixed with 10□μM compound in a buffer containing 20□mM HEPES, pH□7.5, and 500□mM NaCl. SYPRO Orange (5000×, Invitrogen) was added as a fluorescence probe (1 μl per mL). Subsequently, temperature-dependent protein unfolding profiles were measured using the QuantStudio™ 5 realtime PCR machine (Thermo Fisher). Excitation and emission filters were set to 465 nm and 590 nm, respectively. The temperature was raised with a step rate of 3°C per minute. Data points were analyzed with the internal software (Thermal Shift Software™ Version 1.4, Thermo Fisher) using the Boltzmann equation to determine the inflection point of the transition curve.

### Statistics and Reproducibility

All Immunoblot results presented in figures are representative results from at least three independent experiments. All cell death and viability assays (except for the inhibitor screen) were performed in triplicate samples and the data presented are representative of three independent experiments. Inhibitor screen was performed with three biological replicates and the positive hits were reverified in an independent experiment. The calculations, statistical analyses and graphs were performed using Microsoft Excel. Two-tailed unpaired t-test was used to calculate statistical significance for the viability assays. The statistics and source data are presented in Supplementary Table S3.

## Results

### MK2-KO and MK2/3-DKO MEFs predominantly undergo necroptosis rather than apoptosis

Previous studies from us and others have identified MK2-mediated RIPK1 phosphorylation as a checkpoint of receptor mediated RIPK1 activation. TNFα stimulation in the presence of smac mimetics or IKK1/2 inhibitors were shown to sensitize MK2-deficient cells to RIPK1 activation-dependent cytotoxicity. Inactivation of the IKK1/2-dependent checkpoint seems to be upstream to the MK2-dependent checkpoint (Menon et al., 2017; Dondelinger et al., 2017; Jaco et al., 2017). To further understand the mechanism of cell death in MK2/3-deficient cells, we monitored RIPK1 activation and signaling in these cells. Consistent with the previous studies, MK2/3-deficient MEFs displayed higher levels of RIPK1 autophosphorylation (pRIPK1-S166) upon stimulation with TNF and smac-mimetics (SM), compared to the same cells but rescued with retrovirally expressed MK2 (Figure 1A). As expected, the pS166-RIPK1 was further enhanced when the cells were co-treated with the pan-caspase inhibitor zVAD-fmk to induce a switch to necroptosis. Surprisingly, there were significantly higher levels of necroptotic marker phosphorylated and oligomerised MLKL (pS345-MLKL) present in the DKO cells, even in the absence of caspase inhibition. This indicates that MK2/3-deficiency is associated with predominant necroptosis in response to a pro-apoptotic stimulus. Similar results were observed when MK2-deficient MEFs were used instead of MK2/3-DKO cells (Supplementary Figure S1A). In addition to MK2 being a RIPK3 kinases in the cytoplasm, we have also shown strong association between MK2 and RIPK1 in ripoptosome-like complexes in transfected cells (Menon et al., 2017). To rule out any differences between absence of MK2 protein and absence of MK2 activity in the observed phenotype, we performed similar experiments in wild-type (WT) MEFs in the presence and absence of the MK2 inhibitor PF3644022. Interestingly, MK2 inhibition also induced a predominant necroptotic response (Supplementary Figure S1B).

**Figure 1.**
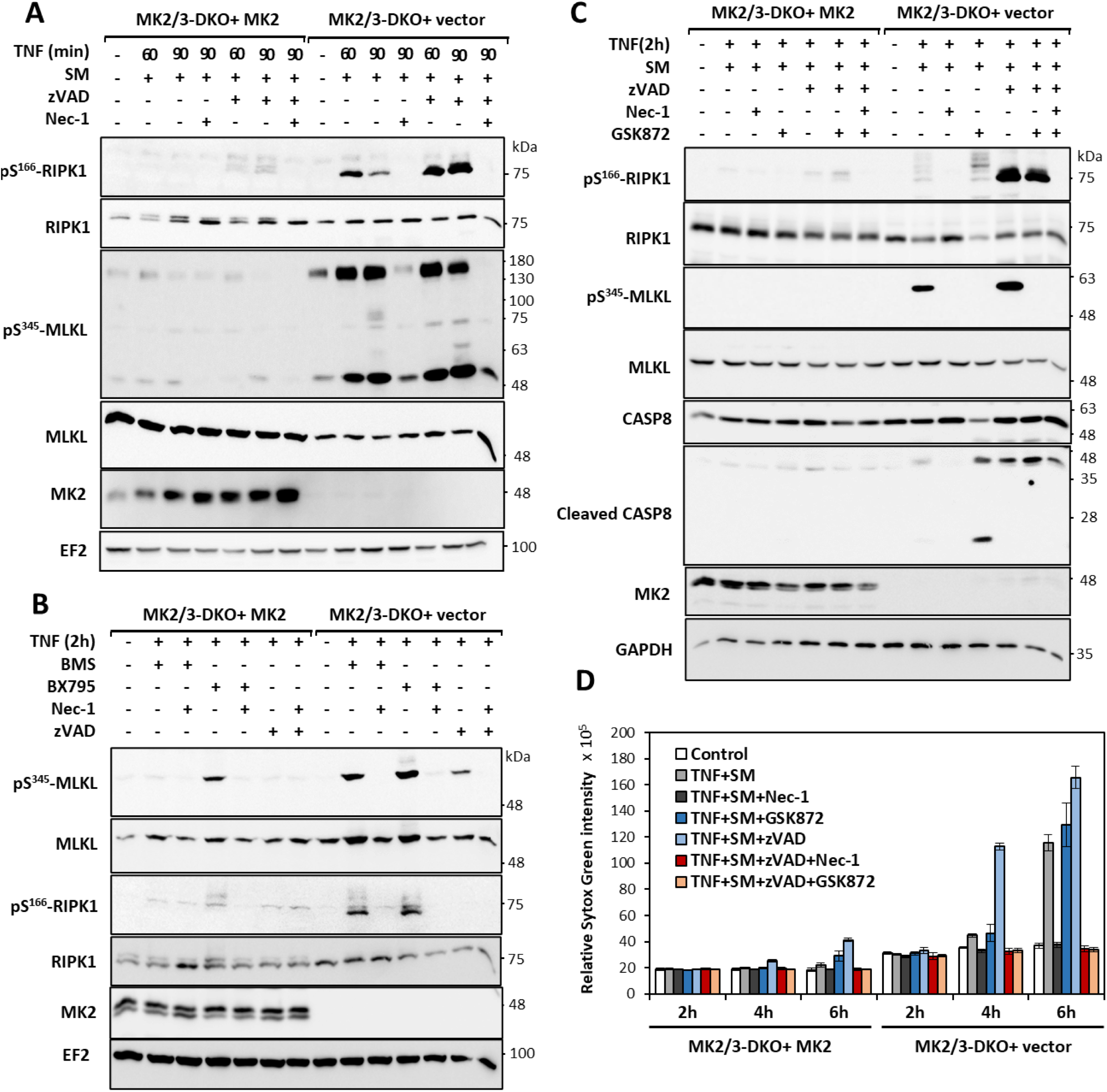
MK2-deficient cells predominantly undergo necroptosis in response to pro-apoptotic stimuli. **A.** MK2/3-deficient MEFs transduced with MK2 expression or control vector are treated with TNF in the presence of smac-mimetics (SM), a pro-apoptotic stimulus, for 60 and 90 minutes. B. Diverse apoptotic stimuli including combinations of TNF with the IKK1/2 inhibitor BMS345541 (BMS) and with BX795 (TBK1 inhibitor) induce predominant necroptotic response in MK2/3-deficient cells. **C.** MK2/3-deficient cells were pre-treated with RIPK3 inhibitor (GSK872) and Smac mimetics (SM) for 30 minutes followed by TNF-treatment for 2 hours. (A-C) Numbers at the right of the blot indicate molecular mass of the marker proteins in kDa. **D.** Sytox-green based cytotoxicity assay was performed with MK2/3-deficient and MK2-rescued cells treated in the presence and absence of caspase inhibitor (zVAD) as indicated.

We then asked the question whether this pro-necroptotic response in the absence of the MK2 checkpoint is specific to SM treatment. When the IKK1/2 inhibitor BMS345541 (BMS) or the TBK1 inhibitor BX795 were combined with TNFα to provide different pro-apoptotic stimuli, we again observed a stronger necroptotic response in the absence of MK2/3, as indicated by RIPK1 and MLKL phosphorylation (Figure 1B). In all these cases, RIPK1 and MLKL phosphorylation were dependent on RIPK1 activity and inhibited by RIPK1 inhibitor necrostatin-1 (Nec-1) (Figure 1A, B and Supplementary Figure S1A, B). It is known that MLKL-S345 phosphorylation is mediated by RIPK3 (Sun et al., 2012). To understand the hyperactivation of RIPK1-RIPK3-MLKL axis in response to the pro-apoptotic TNF-SM stimulus, we pre-treated the cells with the RIPK3 inhibitor GSK872. Interestingly, inhibition of RIPK3 led to significant increase of cleaved caspase-8 (CASP8) in TNF-SM-treated MK2/3-deficient cells, indicative of a switch to apoptosis (Figure 1C). The observations made by immunoblot analyses of necroptotic and apoptotic markers were verified by Sytox-green based cytotoxicity assays, wherein MK2/3-deficiency was associated with strong necrotic death in the presence and absence of caspase inhibitor (Figure 1D). While Nec-1 protected cells independent of the type of cytotoxic stimulus, RIPK3 inhibitor (GSK872) was effective only in combination with the caspase inhibitor (zVAD).

### Screen for small molecules sensitizing MK2 deficient cells to cell death

Sensitivity of MK2-deficient cells to RIPK1-dependent cell death is attributed to the lack of MK2-mediated phosphorylation of RIPK1, which suppresses RIPK1 activation. However, this does not explain the predominant necroptotic death associated with MK2-deficiency. To understand the pathways involved in this process, we aimed identifying additional small molecules which sensitizes MK2-deficient cells to TNF-dependent/ RIPK1-dependent cell death. Therefore, we performed a screen using a library of characterized small molecule kinase inhibitors (Supplementary Table S1). MK2-KO cells transduced with empty vector or rescued with an MK2-expression vector were treated with TNFα alone or together with the inhibitors of the screening panel for 6h and cell viability was quantified by CCK8 colorimetric assay (Figure 2). Consistent with previous findings, TNF-alone was not cytotoxic, however, several small molecules sensitized both MEF lines to TNF (compound 15, 42, 133, 142). More interestingly, CAY10657 (IKK2 inhibitor, comp.14), BIO (GSK3 inhibitor, comp. 32), Bisindolylmaleimide VIII (comp. 54) and IX (comp. 55, PKC inhibitors), PIK-75 (PI3 kinase inhibitor, comp. 94) and 5-Iodo tubercidin (5-ITu, comp.130) displayed a specific sensitization effect in the absence of MK2 (Figure 2 and Supplementary Table S2). The strongest, most significant effect was observed with 5-ITu, an adenosine derivative which acts both as adenosine kinase inhibitor and as protein kinase inhibitor (Massillon et al., 1994; Parkinson et al., 1996; Ugarkar et al., 2000).

**Figure 2.**
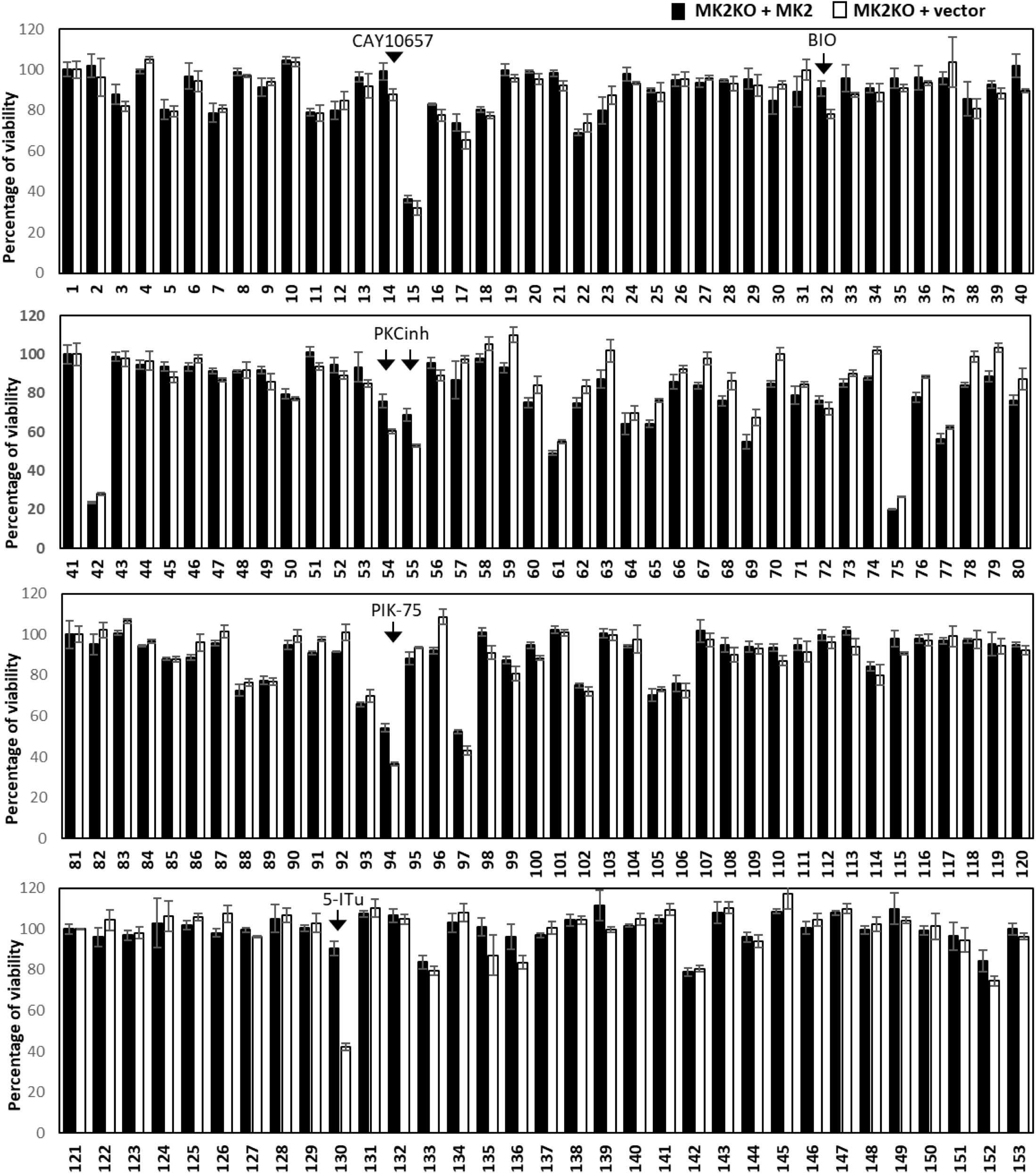
Screen for small molecules facilitating TNF-induced death in MK2-deficient cells. MK2-deficient MEFs transduced with MK2 expression or control vector were treated with TNF alone or in the presence of the inhibitor panel (at 10 μM concentration) for 6h and cell viability was quantified by CCK8 colorimetric assay. Each treatment was performed in triplicates. While TNF alone is not cytotoxic, several small molecules sensitize both MEF lines to TNF (comps. 15, 42, 133, 142). Compound 14/CAY10657 (IKK2 inhibitor), comp. 32/BIO (GSK3 inhibitor), comp. 54,55/Bisindolylmaleimide VIII/IX (PKC inhibitors), comp. 94/PIK-75 (PI3 kinase inhibitor) and comp. 130/5-Iodotubercidin (5-ITu) display specific sensitization effects dependent on MK2-deficiency. Average values of n = 3 independent wells are plotted ± SD.

### 5-ITu sensitizes MK2-KO cells to RIPK1-dependent cell death similar to SM and IKK inhibition

An effect of the adenosine analog 5-ITu on TNF signaling has not been reported so far. To further characterize and confirm the effect of 5-ITu on MK2/RIPK1-mediated cytotoxicity, we analyzed the effect of increasing doses of 5-ITu in sensitizing MK2 KO cells to TNF-induced cytotoxicity. When MK2-KO cells transduced with empty vector control or rescued with MK2 expressing vector were treated with 5-ITu and TNF, dose-dependent loss of viability was observed only in the absence of MK2 (Figure 3A). Similar results were observed when the same experiments were performed with MK2/3-DKO MEFs rescued by MK2 or empty vector (Supplementary Figure S2A). 5-ITU is a nucleoside analog with known cytotoxic effects. To understand whether the MK2-specific effects observed here represent general effects of 5-ITu induced cell death, we monitored the cytotoxic effect of 5-ITu in the absence of TNF. While 5-ITu alone induced general loss of cell viability of MEFs ranging from about 20% after 6 h of treatment to 60% after 24 h treatment, no MK2-deficiency-associated phenotype could be observed (Figure 3B). Even in the MK2/3-DKO background, the presence or absence of MK2 expression had no impact on 5-ITu induced cell death (Supplementary Figure 2B).

**Figure 3.**
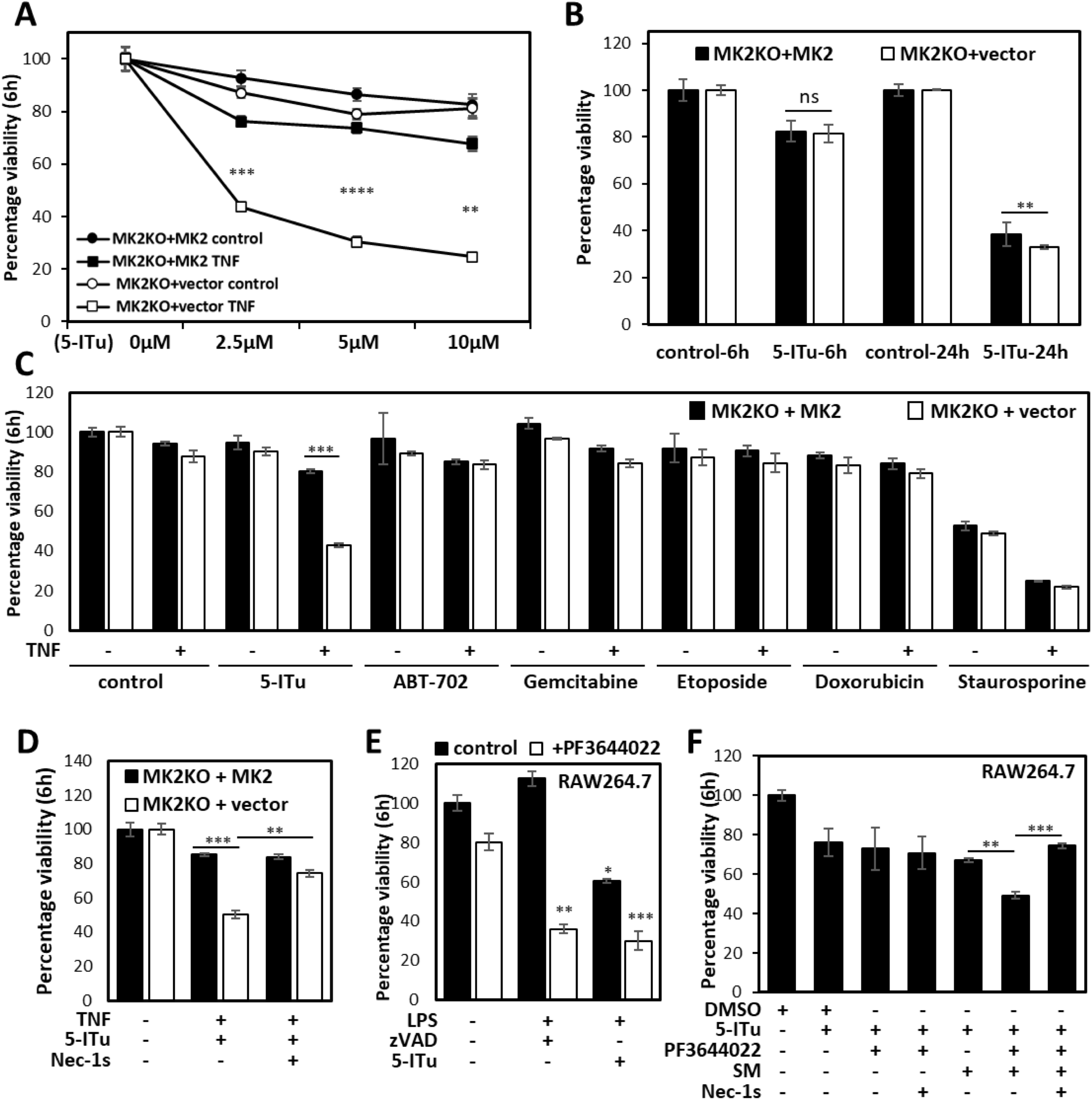
5-Iodotubercidin potentiates RIPK1-dependent cell death in the absence of MK2 activity. **A.** MK2-deficient MEFs transduced with MK2 expression or control vector were treated with different doses of 5-ITu in the presence or absence of TNF (10 ng/mL) for 6h. B. Cells of indicated genotype were treated with 10 μM 5-ITu for 6 and 24h and cell viability was quantified. C. MK2-deficient MEFs transduced with MK2 expression or control vector were treated with indicated small molecules (5 μM ABT702, 100 nM gemcitabine, 5 μM etoposid, 5 μM doxorubicin and 5 μM staurosporine) in the presence or absence of TNF for 6h and cell viability was assessed. D. MK2-deficient MEFs transduced with MK2 expression or control vector were treated as indicated and viability was quantified after 6 h treatment. E, F. RAW264.7 cells were treated as indicated to monitor the effect of 5-ITu in LPS - induced necroptosis (E) and SM-mediated autocrine TNF-dependent death (F). Average values of *n =* 3 independent wells are plotted ± s.d. (** denotes p-value ≤ 0.001, *** denotes p-value ≤ 0.0001, **** denotes p-value ≤ 0.0001).

Since 5-ITu is a nucleoside analog and activator of DNA damage signaling, we decided to compare the effect of another nucleoside analog, gemcitabine, and of other DNA damage-inducing agents, etoposide and doxorubicin, on TNF-induced cytotoxicity. Interestingly and unlike 5-ITu, none of these compounds induced significant cell death alone or in combination with TNF (Figure 3C). We also tested the effect of the pan kinase inhibitor and apoptosis inducer staurosporine on TNF-induced cell death. While staurosporine induced significant loss of cell viability in the presence and absence of TNF, the effects were independent of MK2 expression (Figure 3C). To check whether the specific effect of 5-ITu on TNF-induced cell death is mediated by adenosine kinase inhibition, we used ABT-702 (Radek et al., 2004), a non-nucleoside adenosine kinase inhibitor. Neither MK2-KO controls, nor the MK2-rescued MEFs displayed significant loss of cell viability upon ABT-702 treatment alone or in combination with TNF (Figure 3C). These results indicate that the MK2-specific effects exerted by 5-ITu on TNF-induced cell death is unique to this compound and cannot be explained by general effects on DNA damage signaling, adenosine kinase inhibition or pan-protein kinase inactivation. Similar results were observed when we performed the experiments in the MK2/3 DKO MEFs (Supplementary Figure 2C). Moreover, to monitor the RIPK1-dependence of the TNF-5-ITu-mediated cell death, we treated MK2-KO (Figure 3D) and MK2/3-DKO (Supplementary Figure 2D) MEFs and corresponding cells rescued with MK2 with TNF-5-ITu in the presence of the RIPK1 kinase inhibitor necrostatin-1s (Nec-1s). This RIPK1 inhibitor significantly suppressed TNF-ITu-induced death in MK2 and MK2/3-deficient cells, indicating that the cell death is dependent on RIPK1 activity.

### 5-ITu induces RIPK1-dependent macrophage cell death in combination with LPS and SM

In addition to the TNF receptor, TLR4 is also shown to induce RIPK1-dependent cell death response, which is enhanced in the absence of MK2 activity (Menon et al., 2017). To understand whether the effect of 5-ITu is specific to TNF signaling and MEFs, we monitored the effect of 5-ITu in LPS-induced necroptosis of RAW264.7 macrophages. When RAW264.7 cells were treated with the previously characterized necroptotic stimulus LPS-zVAD for 6 h, significant death was observed only in the presence of the MK2 inhibitor PF3644022 (Figure 3E). Combined LPS-5-ITu-stimulation also led to significant loss of cell viability even in the absence of PF3644022, but the cytotoxicity was significantly enhanced by MK2 inhibition using PF364402 (Figure 3E). We then asked the question whether 5-ITu may have any additive/synergistic effect with the SM-mediated autocrine TNF-dependent death in macrophages. Interestingly, we observed a similar MK2 inhibitor-dependent sensitizing effect of 5-ITu on the cell death induced by SM, which was rescued by the RIPK1 inhibitor Nec-1s (Figure 3F). Again, 5-ITu alone displayed some reduction in the viability of cells after 6 h, which was neither MK2-, nor RIPK1-dependent.

### 5-ITu sensitizes cells to RIPK1-dependent necroptosis by downregulating IKK signaling

To understand the mechanism by which 5-ITu sensitizes MK2-deficient cells to RIPK1-dependent cell death, we compared TNF-induced cell death signaling in the presence of IKK1/2 inhibitor BMS with such signaling in the presence of 5-ITu. Interestingly, both BMS345541 and 5-ITu induced RIPK1 activation and downstream necroptotic MLKL phosphorylation in MK2-deficient cells (Figure 4A). This indicates, similarly to SM and BMS345541, that 5-ITu-mediated cytotoxicity in the absence of MK2 activity is predominantly necroptotic. Then we performed a detailed investigation of signaling in the MK2/3-DKO MEFs transduced with empty or MK2 expression vector. Consistent with previous experiments, there was prominent RIPK1 activation (pS166-RIPK1) and MLKL phosphorylation (pS345-MLKL) in the absence of MK2 activity (Figure 4B). At the 3h-stimulation time-point analyzed, there was no detectable upregulation of the three canonical MAPK pathways with TNF alone and pERK1/2, pJNK and pp38 signals were mostly downregulated in stimulated cells. A20/TNFAIP3, a ubiquitin editing enzyme and NFκB feedback regulator was upregulated by TNF, downregulated by 5-ITu treatment and stayed low in response to all the cytotoxicity inducing combinatory stimuli (Figure 4B). The late-stage MAPK activation was predominantly downstream to necroptotic signaling as the activation corresponded to pMLKL and was inhibited by RIPK1 and RIPK3 inhibitors. Interestingly, 5-ITu treatment led to the complete loss of IκBα protein, independent of TNF stimulation and RIPK1 activation, suggesting this to be a direct consequence of 5-ITu (Figure 4B).

**Figure 4.**
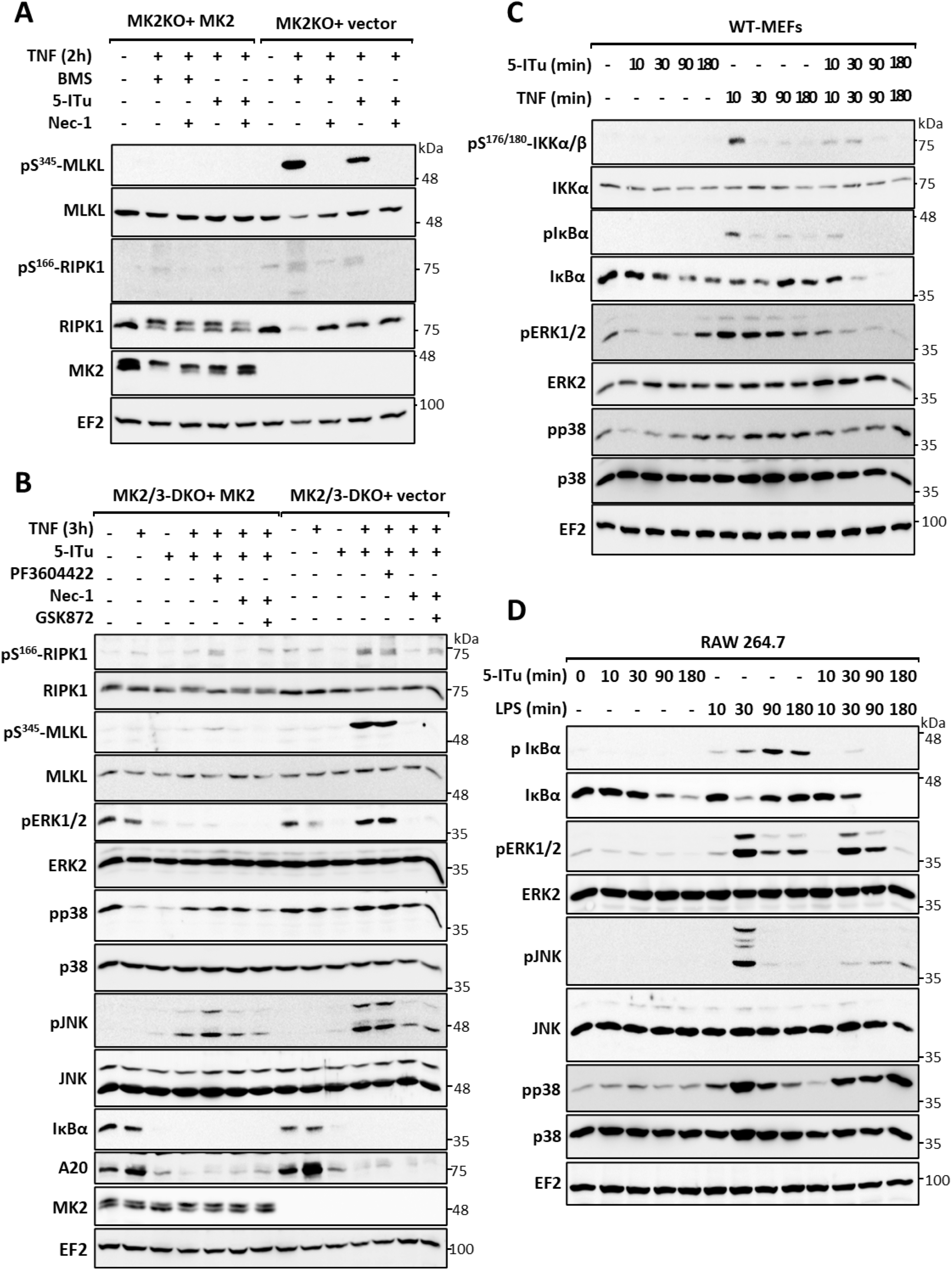
5-ITu sensitizes cells to RIPK1-dependent necroptosis by downregulating IKK signaling. **A.** MK2-deficient MEFs transduced with MK2 expression or control vector were treated with IKK1/2 inhibitor BMS and 5-ITu in combination with TNF for 2h. MLKL and RIPK1 phosphorylation was monitored with additional control blots. RIPK1 band shift induced by MK2-mediated phosphorylation is absent in MK2-KO cells and EF2 is shown as loading control. **B.** MK2/3-deficient MEFs transduced with MK2 expression or control vector were treated with 5-ITu alone or in combination with TNF for 3h. Cells were pre-treated for 30 minutes with the MK2 inhibitor PF3604422, the RIPK1 inhibitor Nec-1 or the RIPK3 inhibitor GSK872, as indicated. **C.** Wild type (WT) - MEFs were treated with 5-ITu or TNF alone or in combination for a short-term kinetics (10-180 minutes). **D.** RAW264.7 cells were treated with 5-ITu or LPS alone or in combination for 10-180 minutes and signaling was monitored by immunoblotting as indicated. (A-D) Numbers at the right of the blot indicate molecular mass of the marker proteins in kDa.

To analyze the effect of 5-ITu on NFkB-signaling in more detail, we performed short-term kinetics of 5-ITu, TNF and 5-ITu together with TNF in wild-type (WT) MEFs. As expected, TNF stimulation led to a transient increase in IKK1/2 activation and IκBα phosphorylation, which was blocked by 5-ITu (Figure 4C). While 5-ITu led to a time-dependent decrease in IκBα levels, TNF-treatment led to a transient decrease at 10-30 minutes, followed by increase probably owing to its re-synthesis. When cells were treated with a combination of 5-ITu and TNF, there was a complete loss of IκBα re-synthesis, which is a mark of NFκB pathway inhibition (Figure 4C). These data clearly indicate that 5-ITu-mediated effects on TNF signaling were caused by suppression of NFκB activity. To test whether this is also true for the case of LPS-mediated effects, we performed additional experiments in RAW264.7 macrophages. Thirty minutes of LPS stimulation resulted in activation of all three MAPK, with JNK phosphorylation being more transient than pERK1/2 and pp38. While co-treatment with 5-ITu led to the suppression of JNK and ERK1/2 phosphorylation, a sustained p38 phosphorylation was detected when compared to LPS alone (Figure 4D). Similarly, as observed in MEFs, 5-ITu led to a time-dependent decrease in IκBα levels also in macrophages. LPS-stimulation induced strong and sustained IκBα phosphorylation, while there was no detectable phosphorylation in cells treated with 5-ITu alone. LPS induced degradation of IκBα was detectable at 30 minutes with clear re-synthesis of this protein visible at later time-points. Similar to the data for MEFs, 5-ITu strongly suppressed IκBα re-synthesis (Figure 4D). Thus, 5-ITu seems to impair both TNF- and LPS-mediated signaling by interfering with NFκB pathway activity.

## Discussion

We and others have previously shown that MK2 limits RIPK1 activation by phosphorylating it at S320/S335 residues (S321/336 in murine RIPK1) in response to pro-inflammatory stimuli such as TNF, LPS and *Yersinia* infection, and therefore, can control RIPK1-dependent cell death (Menon et al., 2017). While earlier reports have not specifically looked at the mode of cell death facilitated by the loss of MK2, our data obtained here clearly demonstrate that, even under pro-apoptotic conditions, MK2-deficient cells predominantly die by necroptosis. This is unusual, since induction of necroptosis as a primary cytotoxic response to TNF/LPS *in vitro* usually require CASP8 inhibition or FADD depletion. However, it has been established that CASP8 inhibition is not a prerequisite for necroptotic death *in vivo* (Duprez et al., 2011). Previously, proteasome inhibition was shown to induce necroptosis in MEFs and human leukemia cells without the need of caspase inhibition and TNF stimulation, but through the accumulation of K48-poly-ubiquitinated RIPK3 (Moriwaki et al., 2016). In this case, cell death was not RIPK1 activity-dependent, but RIPK3 inhibition switched it to apoptosis, as observed in the case of MK2-deficiency. It has been recently shown that the accumulation of a critical amount of active MLKL at the plasma membrane is crucial for execution of necroptosis (Samson et al., 2020). Therefore, it is also possible that the level of active RIPK1 available to activate necroptotic cascade is a crucial parameter for deciding cell death outcome. We further noted that inhibition of RIPK1 activity completely abrogated TNF-induced cell death in MK2-deficient cells treated with pro-apoptotic stimuli, while inhibition of RIPK3 activity predominantly resulted in apoptotic cell death. This is consistent with a previous study showing that inhibition of RIPK3 blocks necroptosis but induces apoptosis (Mandal et al., 2014). Interestingly, expression of a RIPK1 deletion mutant lacking the intermediate domain which harbors the MK2 phosphorylation sites was shown to induce a shift from TNF-induced necroptosis to apoptosis in murine L929 cells (Duprez et al., 2012). The prominent caspase cleavage site on RIPK1 (D324) is also part of the intermediate domain and very close to the MK2 target site S320. The finding that mutations, which render RIPK1 caspase-resistant, are associated with autoinflammatory disease (Lalaoui et al., 2020), further supports a role for this region of the kinase and the post-translational modifications therein for cell-fate and apoptosis/necroptosis decisions by RIPK1. Apart from this, loss of the p38/MK2 upstream kinase TAK1 was shown was associated with increased sensitivity to TRAIL induced necroptosis without the need for caspase inhibition (Goodall et al., 2016). This study revealed a novel connection between autophagosomes and necroptosis and showed RIPK1 activation and necrosome assembly templated by autophagosomes. In this case, depletion of autophagy machinery components, induced the necroptosis – apoptosis switch (Goodall et al., 2016).

The screen for molecules which sensitize MK2-KO MEFs to TNF-induced necroptosis revealed 5-ITu as the most prominent hit. The data indicated that 5-ITu, but not related pan-kinase inhibitors or adenosine kinase inhibitors, displayed this novel MK2-deficiency-associated phenotype of cells death. Moreover, we also verified a similar sensitizing effect of 5-ITu on the LPS-induced necroptotic response in macrophages. These findings indicated that 5-ITu was attenuating a RIPK1 activation checkpoint distinct from the MK2-dependent checkpoint, which has an additive effect with inactivation of p38/MK2 pathway. These findings are reminiscent to data on other RIPK1 checkpoint inhibitors of TBK1 and IKKs signaling. Consistently, we have observed significant effect of 5-ITu in basal and TNF/LPS-induced NFκB signaling, which was mainly evident in cells with defective IκBα resynthesis. While originally discovered as an adenosine kinase inhibitor, with some inhibition on protein kinases CK1 and CK2, 5-ITu was later shown to strongly inhibit ERK2 (Fox et al., 1998). Interestingly, a subsequent study used 5-ITu as an ERK1/2 inhibitor demonstrating inhibitory effects on silica-induced phosphorylation of p55 TNF receptor in RAW264.7 macrophages. In this study, 5-ITu also enhanced silica-induced macrophage apoptosis, a phenotype which was also promoted by dominant negative IκBα mutants (Gambelli et al., 2004). Based on our findings, these data need to be re-evaluated to learn whether these effects can also be attributed to 5-ITu-mediated inhibition of NFκB pathway, too. In addition to ERK2, Haspin/GSG2 is another well-characterized target of 5-ITu (Antoni et al., 2012; Karanika et al., 2020; Heroven et al., 2018). In order to identify the direct target(s) of 5-ITu relevant to TNF-induced RIPK1 checkpoint signaling, we performed a differential scanning fluorimetry (DSF)-based selectivity kinome screen using about 100 protein kinases. The screen identified a number of kinase targets of 5-ITu which showed similar temperature shift upon 5-ITu binding comparable to the reference compound staurosporine, indicating strong inhibition. We identified about 20 kinases which displayed more than 3-degree shift in Tm, with the top positions with more than 10 degrees shift occupied by CLK1 (13.3), GSG2/Haspin (12.3) and DYRK2 (10.8) (Supplementary Figure 3A and Supplementary Table S3). A CLK1/4 inhibitor was also part of our original screen (TG003, Supplementary Table 2) and did not show any effect comparable to 5-ITu. Death assays performed in the presence of LDN192960 (DYRK2 and Haspin inhibitor) and T3-CLK (CLK1/2/3 inhibitor) also did not display any effect on TNF-induced cell death in the MK2-KO cells (data not shown). This leaves the explanation open that the phenotype on cell death and NFκB pathway inhibition shown by 5-ITu could be a combinatorial effect of inhibition of more than one protein kinase or a so far undiscovered target of 5-ITu. Interestingly, when we used PathwayNet tool (Park et al., 2015) (https://pathwaynet.princeton.edu/) to understand functional interactions between the top candidates from the screen (≥5 degree ΔTm, n=13), it revealed a high confidence, closely linked functional network between these protein kinases indicating a possible cooperation of these enzymes also *in vivo* (Supplementary Figure 3B).

Cell death control in development, inflammation and infection is coordinated by RIPK1, but only a small part of the complex post-translational modification (PTM) code of RIPK1 is understood so far. In parallel to the first decoding of its PTMs, RIPK1-dependent necroptosis has become a sought-after target for positive and negative interference against cancer, neurodegeneration and inflammatory pathologies (Degterev et al., 2019). The identification of 5-ITu - an FDA-approved compound and genotoxic drug with anti-cancer potential - as modulator of RIPK1-dependent death, in combination with p38/MK2 inhibitors and smac mimetics opens possibilities for future therapies. Moreover, this adds 5-ITu to the list of anti-cancer agents such as Dabrafenib, Vemurafenib, Sorafenib, Pazopanib and Ponatinib, which are all multi-kinase targeting inhibitors and display therapeutic potential in diverse pathologies by interfering with the necroptosis pathway (Fulda, 2018).

## Conflict of Interest

The authors declare no conflicts of interests.

## Acknowledgements

This work was supported by the Deutsche Forschungsgemeinschaft (DFG) grants ME4319/3-1 (M.B.M.) and GA453/16-1 (M.G.). SK and AK are grateful for support by the the SGC, a registered charity (number 1097737) that receives funds from the Bayer Pharma AG, Boehringer Ingelheim, Canada Foundation for Innovation, Eshelman Institute for Innovation, Genome Canada, Innovative Medicines Initiative (EU/EFPIA) [ULTRA-DD grant no. 115766], Janssen, Merck KGaA Germany, MSD, Novartis Pharma AG, Ontario Ministry of Economic Development and Innovation, Pfizer, Takeda, and the Frankfurt Cancer Institute (FCI) and the DKTK cancer consortium.

## Author Contribution Statement

C.C. performed majority of the experiments, designed experiments, and analysed data. A.Kr. and S.K. designed and performed the DSF experiment. M.W. provided protocols and intellectual input. A.Ko. and M.G. designed experiments and gave conceptual insights. M.G. and M.B.M procured funding. M.B.M. designed and supervised experiments and analyzed data. M.G., C.C. and M.B.M. prepared the manuscript.

## Data Availability Statement

All data generated during this study leading to the findings presented here are included in this published article and its supplementary data files.

## Notes

### Competing Interest Statement

The authors have declared no competing interest.

### Summary of Updates

- Amentments in the main text - Revised Suppl. Table S3

