## Supplementary Figures 1-3 for "5-iodotubercidin sensitizes cells to RIPK1-dependent necroptosis by interfering with NFκB signaling"

### Slide 1
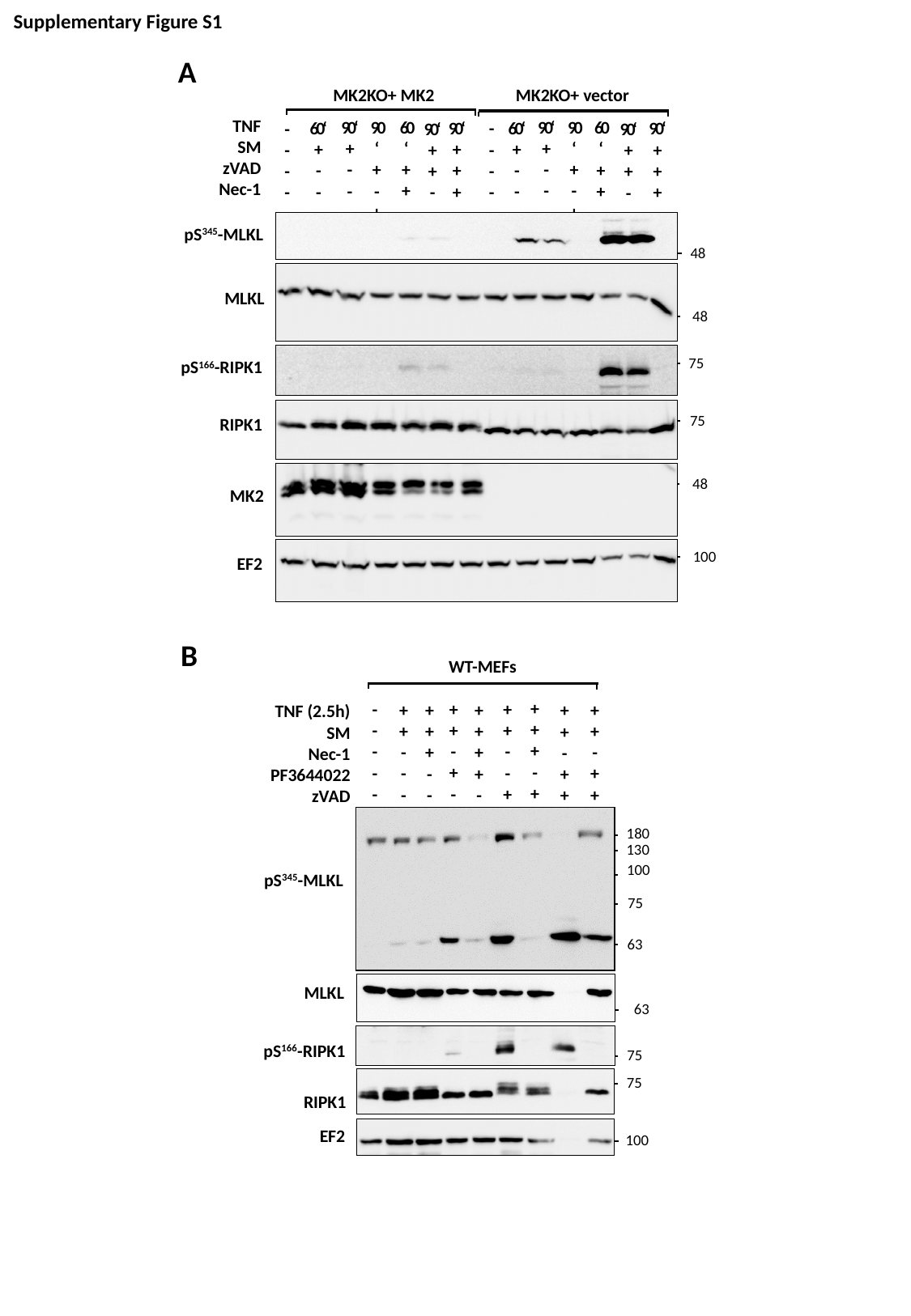

Supplementary Figure S1
A
MK2KO+ MK2
MK2KO+ vector
TNF
SM
zVAD
Nec-1
60‘
+
+
-
90‘
+
-
-
90‘
+
-
+
60‘
+
-
-
90‘
+
+
+
-
-
-
-
90‘
+
-
-
90‘
+
-
+
60‘
+
+
-
60‘
+
-
-
90‘
+
+
+
-
-
-
-
90‘
+
+
-
90‘
+
+
-
48
pS^345^-MLKL
48
MLKL
75
pS^166^-RIPK1
75
RIPK1
48
100
MK2
EF2
B
WT-MEFs
+
+
+
-
+
+
+
-
+
-
+
+
-
-
+
-
-
-
-
-
+
+
-
+
+
+
+
-
-
-
+
+
+
-
-
+
+
+
+
-
+
+
-
+
+
TNF (2.5h)
SM
Nec-1
PF3644022
zVAD
180
130
100
75
63
pS^345^-MLKL
63
MLKL
75
pS^166^-RIPK1
75
RIPK1
EF2
100

### Slide 2
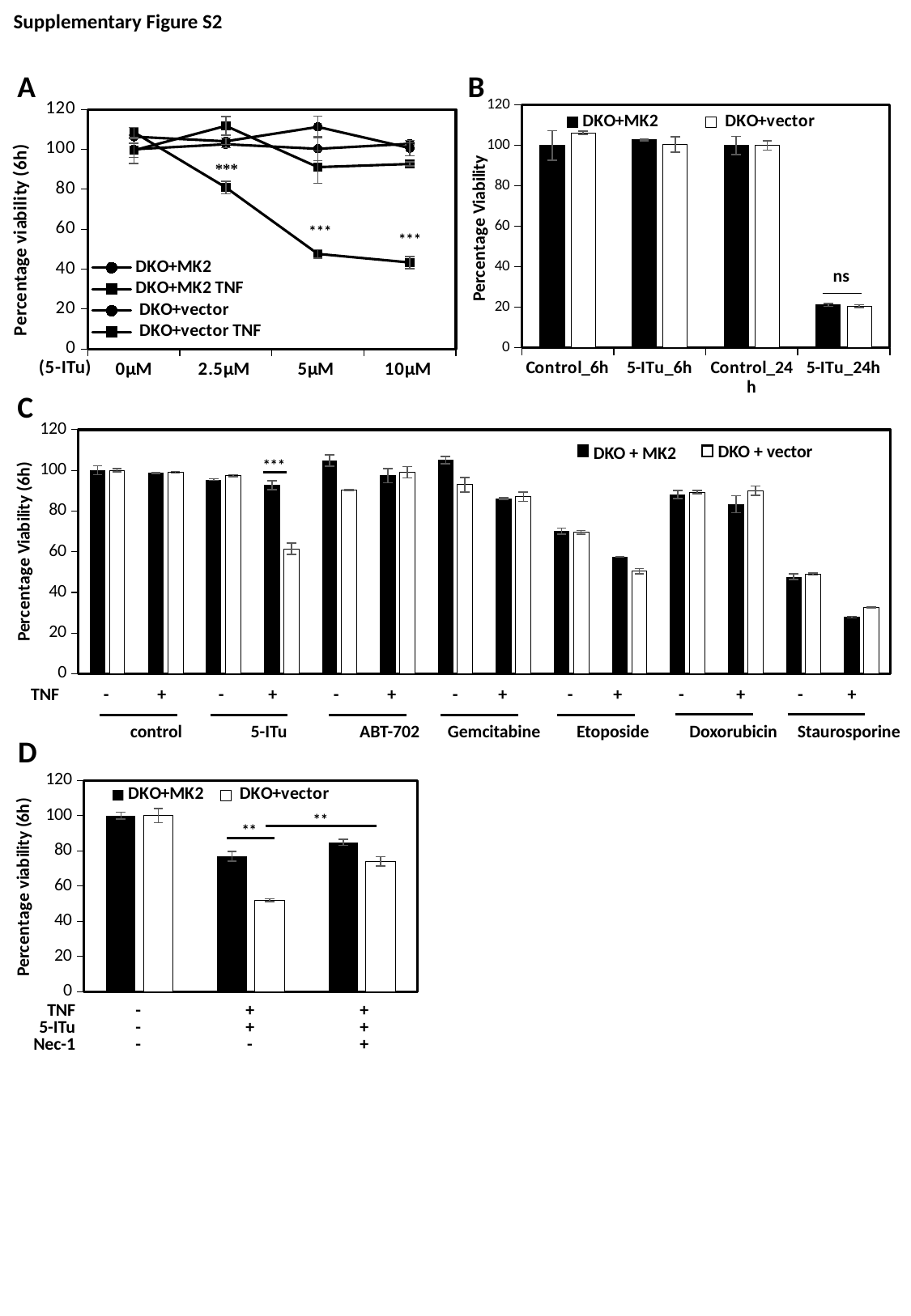

Supplementary Figure S2
#### Chart
| Category | DKO+MK2 | DKO+vector |
|---|---|---|
| Control_6h | 100.0 | 106.2916151670201 |
| 5-ITu_6h | 102.73563611121388 | 100.58253777494632 |
| Control_24h | 100.0 | 100.0 |
| 5-ITu_24h | 21.134164874579216 | 20.48202479037535 |A
B
#### Chart
| Category | DKO+MK2 | DKO+MK2 | DKO+vector | DKO+vector |
|---|---|---|---|---|
| 0µM | 100.0 | 99.46862112367818 | 106.2916151670201 | 108.53012925470695 |
| 2.5μM | 102.5154462509755 | 111.67878864695263 | 103.97927182152955 | 80.88213428673252 |
| 5μM | 100.24366686411646 | 91.04971783962584 | 111.1833082088245 | 47.63090924895993 |
| 10μM | 102.73563611121388 | 92.61881482002282 | 100.58253777494632 | 43.33032723367544 |ns
C
#### Chart
| Category | - TNF | - TNF | | +TNF | +TNF |
|---|---|---|---|---|---|
| Control | 100.0 | 100.0 | None | 98.61919282604077 | 99.08859940377162 |
| 5-Itu | 95.5104050227444 | 97.34171307749664 | None | 92.72466183850445 | 61.45689016393845 |
| ABT702 | 104.88366833594546 | 90.22997247499698 | None | 97.4311149143084 | 99.01019248147327 |
| Gemcitabine | 105.08351439278597 | 92.98244552974256 | None | 86.11658462871765 | 86.94981655991158 |
| Etoposide | 70.09115432338014 | 69.5128465798813 | None | 57.425283412812114 | 50.391400599240036 |
| Doxorubicin | 88.23152795783653 | 89.31399030655722 | None | 83.32420417206764 | 90.03090313657447 |
| Staurosporine | 47.715993380054556 | 49.070045330078365 | None | 27.767567031631703 | 32.49125106310111 |DKO + vector
DKO + MK2
***
TNF - + - + - + - + - + - + - +
 control 5-ITu ABT-702 Gemcitabine Etoposide Doxorubicin Staurosporine
D
#### Chart
| Category | DKO+MK2 | DKO+vector |
|---|---|---|
| Control | 99.99999999999999 | 100.00000000000001 |
| TNF+ITu | 76.94662916555664 | 52.02002586472258 |
| TNF+ITu+Nec1 | 84.79028331784845 | 74.02680631985697 |**
**
TNF
5-ITu
Nec-1
-
-
-
+
+
-
+
+
+

### Slide 3
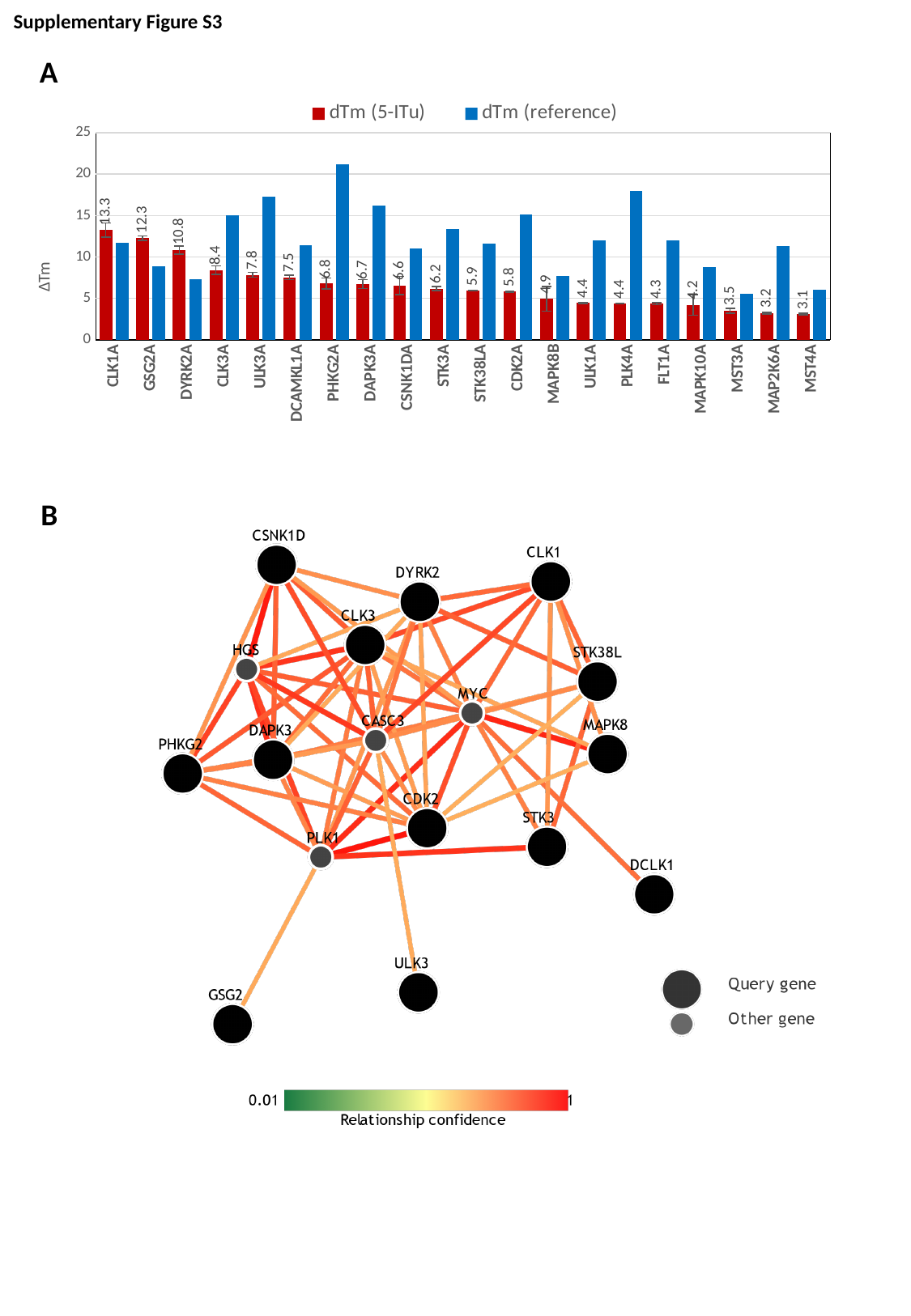

Supplementary Figure S3
A
#### Chart
| Category | | |
|---|---|---|
| CLK1A | 13.261778 | 11.7 |
| GSG2A | 12.268427 | 8.9 |
| DYRK2A | 10.846197 | 7.3 |
| CLK3A | 8.411224 | 15.0 |
| ULK3A | 7.831053 | 17.3 |
| DCAMKL1A | 7.5397415 | 11.4 |
| PHKG2A | 6.7872753 | 21.2 |
| DAPK3A | 6.71373 | 16.246666666666666 |
| CSNK1DA | 6.5674553 | 11.0 |
| STK3A | 6.1773186 | 13.4 |
| STK38LA | 5.9174232 | 11.613333333333332 |
| CDK2A | 5.804573 | 15.166666666666666 |
| MAPK8B | 4.931576 | 7.7 |
| ULK1A | 4.441231 | 12.0 |
| PLK4A | 4.359617 | 18.0 |
| FLT1A | 4.3407726 | 12.0 |
| MAPK10A | 4.202055 | 8.8 |
| MST3A | 3.4949837 | 5.5 |
| MAP2K6A | 3.213169 | 11.339999999999998 |
| MST4A | 3.0983543 | 6.0 |B
