## Supplementary material for "5-iodotubercidin sensitizes cells to RIPK1-dependent necroptosis by interfering with NFκB signaling": Legends to Supplementary Information

**Supplementary Information Legends**

1. **Supplementary Figures S1-S3 (single .pdf file)**

**Supplementary Figure 1. MEFs predominantly undergo necroptosis rather than apoptosis upon MK2 inactivation** **A.** MK2 deficient MEFs transduced with MK2 expression vector or control vector are treated with TNF+Smac-mimetics (SM), a pro-apoptotic stimulus, for 60 and 90 minutes and cell death signaling was monitored by pMLKL and pRIPK1 immunoblotting. MLKL, RIPK1, MK2 and EF2 blots are shown as controls. **B.** Wild Type MEFs were treated with TNF+SM in the presence or absence of MK2 inhibitor PF3644022 for 2.5 hours and analysed similarly as in panel A.

**Supplementary Figure 2. 5-ITu potentiates RIPK1-dependent cell death in the absence of MK2 activity A.** MK2/3 deficient MEFs transduced with MK2 expression vector or control vector were treated with different doses of 5-ITu in the presence or absence of TNF (10 ng/mL) for 6h. **B.** Cells of indicated genotypes were treated with 10 µM 5-ITu for 6 and 24h and cell viability was quantified and represented. **C.** MK2/3 deficient MEFs transduced with MK2 expression vector or control vector were treated with indicated small molecules ( 5 µM ABT702, 100 nM gemcitabine, 5 µM etoposide, 5 µM doxorubicin and 5 µM staurosporine) in the presence or absence of TNF for 6h and cell viability was assessed. **D.** MK2/3 deficient MEFs transduced with MK2 expression vector or control vector were treated as indicated and viability quantified after 6 h treatment. Average values of *n* = 3 independent wells are plotted ± s.d. (** denotes p-value ≤ 0.001, *** denotes p-value ≤ 0.0001, **** denotes p-value ≤ 0.0001).

**Supplementary Figure 3. Differential Scanning Fluorimetry screen for 5-ITu target kinase identification. A.** A DSF assay was performed as detailed in the methods section and the candidates which showed a delta-Tm (fluorescence shift) >3 degrees are plotted with the reference shift shown in parallel (mean +/- standard error plotted). Details of the reference is available in Supplementary Table S3. B. The top 13 candidates from the screen with ≥5-degree shift was used to generate a functional interaction network using the PathwayNet tool (<https://pathwaynet.princeton.edu/>). The minimum relationship confidence cut-off was high at 66%. With the help of additional four genes/proteins, the potential 5-ITu targets form a closely linked network of functional linkages.

2. **Supplementary Table S1. Small molecule library used for cell viability screen.** Names, potential targets and additional details of the kinase inhibitor library used for screening (MS Excel file).
3. **Supplementary Table S2. Source data for small molecule inhibitor screen in Figure 2**. Excel file summarize the results of the cell viability assay screen. The data for 149 compounds treated with 4 sets of DMSO control wells, performed in triplicates are presented as mean with standard deviation (MS Excel file).
4. **Supplementary Table S3. Differential Scanning Fluorimetry based profiling of 5-ITu specificity for kinases.** The raw values from the DSF assay results presented in Supplementary Figure 3 showing the mean fluorescence shift, standard error and the reference shift and identity of the reference inhibitor used in each case (MS Excel file).
5. **Supplementary Table S4: Statistics and source data for the quantitative results.** The actual values (source data) and the *p-*values calculated for the quantitative cell viability data presented in the manuscript (MS Excel file)
